# The anti-immune dengue subgenomic flaviviral RNA is found in vesicles in mosquito saliva and associated with increased infectivity

**DOI:** 10.1101/2021.03.02.433543

**Authors:** Shih-Chia Yeh, Wei-Lian Tan, Avisha Chowdhury, Vanessa Chuo, R. Manjunatha Kini, Julien Pompon, Mariano A. Garcia-Blanco

## Abstract

Mosquito transmission of dengue viruses to humans starts with infection of skin resident cells at the biting site. There is great interest in identifying transmission-enhancing factors in mosquito saliva in order to counteract them. Here we report the discovery of high levels of subgenomic flaviviral RNA (sfRNA) in dengue virus 2-infected mosquito saliva. We show that salivary sfRNA is protected in detergent-sensitive, protease-resistant compartments. Furthermore, we show that incubation with mosquito saliva containing higher sfRNA levels results in higher virus infectivity in human cells. Since sfRNA potently inhibits innate immunity in human cells, we posit that sfRNA in mosquito saliva is present in extracellular vesicles that deliver it to cells at the biting site to inhibit innate immunity and enhance dengue virus transmission.

## Introduction

Half of the world’s human population is at risk of infection with dengue viruses (DENV) transmitted by *Aedes aegypti* mosquitoes (Pierson and Diamond, 2020). Since DENV have evolved to harness mosquito biology to enhance transmission to humans, understanding the transmission cycle of DENV is important for controlling the spread of this disease. Recently, it has become clear that mosquito saliva enhances infection in human cells (Wichit et al., 2016), mouse models (Cox et al., 2012), and non-human primates (McCracken et al., 2020). The mechanistic underpinings for this enhancement are beginning to be understood (Jin et al., 2018; Sun et al., 2020), and here we present evidence for a novel potential transmission enhancer, the subgenomic flaviviral RNA (sfRNA).

sfRNA is a non-coding RNA (ncRNA) produced from partial degradation of the flaviviral RNA genome (gRNA) by host 5’-3’ exoribonucleases that stall at secondary structures in the 3’ untranslated region (UTR) (Pijlman et al., 2008). Data from many groups demonstrate that sfRNA downregulates immune responses in both human and insect cells (Bidet and Garcia-Blanco, 2014; Slonchak and Khromykh, 2018). For instance, DENV serotype 2 (DENV2) sfRNA inhibits the innate immune response by limiting interferon (IFN) expression (Manokaran et al., 2015) and translation of IFN stimulated gene products (Bidet et al., 2014), promoting DENV2 propagation in human cells. Moreover, Zika virus (ZIKV) sfRNA can modulate mRNA decay and splicing and likely limit the effective response of cells to viral infection (Michalski et al., 2019). In DENV2-infected mosquitoes sfRNA is expressed at high levels in salivary glands and downregulates expression of Toll pathway-associated genes *Rel1a* and *CecG* (Pompon et al., 2017). In mosquitoes infected with ZIKV, sfRNA suppresses apoptosis in mosquito tissues and this enhances virus dissemination and accumulation in saliva (Slonchak et al., 2020). Strong evidence of the epidemiological function of sfRNA comes from studies that show that DENV2 strains with higher sfRNA levels, in both human and mosquito, demonstrate higher epidemiological fitness (Bennett et al., 2006; Manokaran et al., 2015; Pompon et al., 2017). Here, we discover that sfRNA is secreted into the saliva of DENV2-infected mosquitoes and is found in detergent-sensitive structures that we propose are extracellular vesicles capable of transmitting the sfRNA to cells at the biting site in human skin. Importantly, our functional data reveal that levels of secreted sfRNA positively correlate with efficiency of DENV2 replication in human cells.

## Results and Discussion

There are several species of DENV2 sfRNA produced by differential exonuclease stalling in mosquito cells and mosquitoes (Filomatori et al., 2017). In *Ae. aegypti* orally-infected with DENV2 NGC strain, we detected three of the expected species, sfRNA1-3, and one decay intermediate, sfRNAx (Fig. 1A). Since sfRNA1 was the major species detected, we quantified the sfRNA1 copies using RT-qPCR as previously described (Pompon et al., 2017). At 10 days post-oral infection (d.p.i.), gRNA was present in salivary glands and in saliva (geometric mean of 3.6 × 10^4^ copies per infected saliva expectorated by one mosquito, which we refer to as saliva sample) (Fig. 1B). We detected sfRNA1 in 100% of infected salivary glands and, importantly, in 91% of infected saliva samples (geometric mean of 1.1 × 10^7^ sfRNA copies per infected saliva sample; 95% CI; lower = 5.6 × 10^6^; upper =2.2 × 10^7^) (Fig. 1C). To normalize sfRNA to its precursor and infection level (estimated by the copies of gRNA), we calculated the ratio of sfRNA: gRNA. The ratio in salivary glands was 720 (± 221.5) and not significantly different (t-test; p = 0.47) from the ratio in saliva, which was 559.2 (± 89.07) (Fig. 1D). We also detected sfRNA in saliva from mosquitoes orally-infected with three other DENV2 strains: PR1940, PR6452, and PR9963 (Figure 1, supplement 3). To validate the presence of sfRNA in infected saliva with an independent approach, we used the well-established method of RNA circularization and sequencing used by Filomatori et al to determine the sequence of sfRNA species (Filomatori et al., 2017). We detected sfRNA1 and sfRNA3 sequences in DENV2 infected mosquitoes (Fig. 1, supplement 4) and in saliva (Fig. 1E). These results led us to conclude that the potent anti-immune sfRNA is found in the saliva of DENV2-infected mosquitoes.

**Figure 1.**
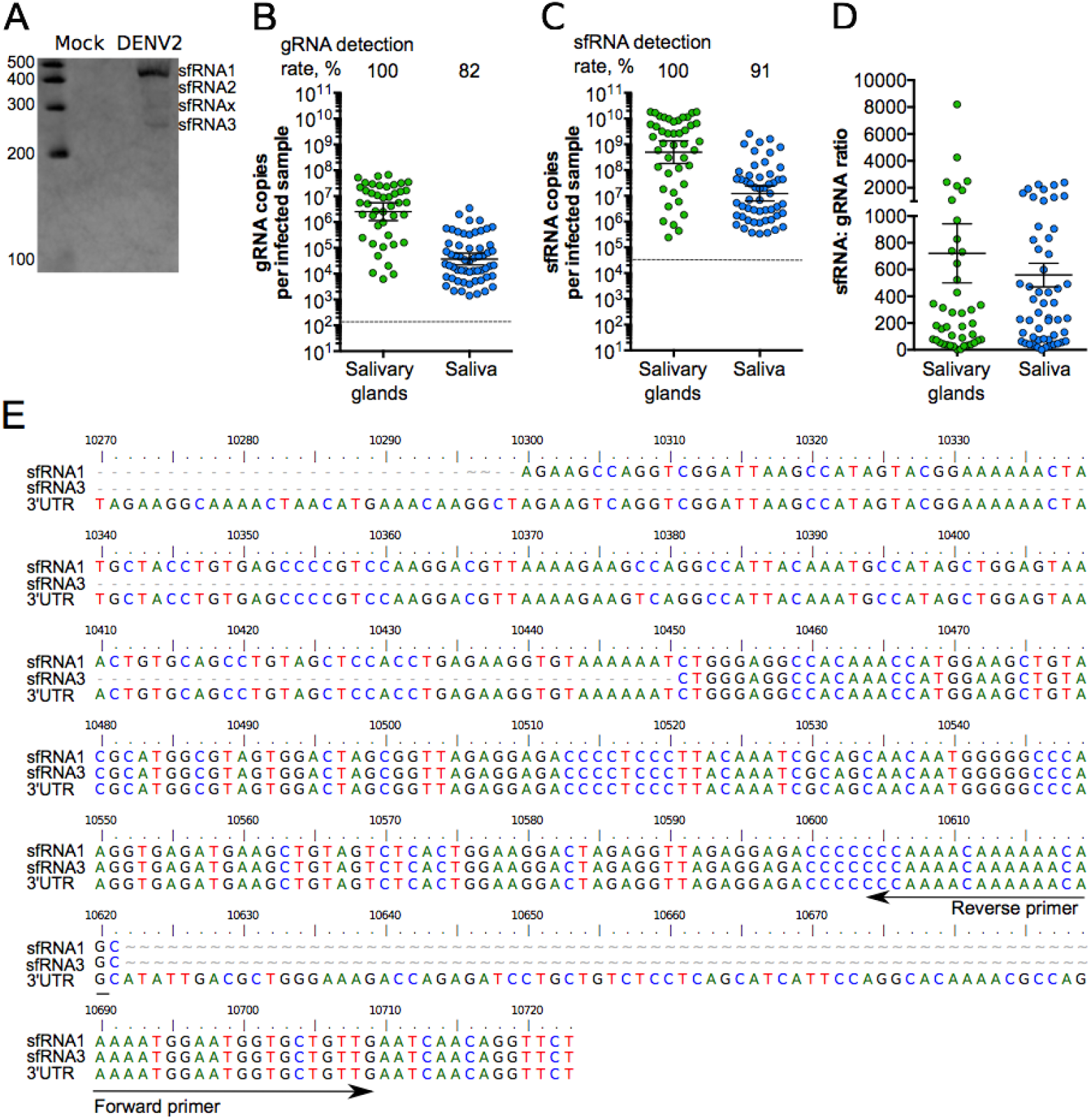
Detection of sfRNA in mosquito saliva. (**A**) sfRNA species detected by Northern blotting in infected mosquitoes. (**B**) gRNA; (**C**) sfRNA; and (**D**) sfRNA:gRNA ratio in salivary glands and saliva. Points represent a pair of salivary glands or one saliva collected from one mosquito. (**B**-**C**) Bars represent geometric means ± 95% C.I. and dashed lines indicate limit of detections which were 133 copies in gRNA (**B**) and 31,216 copies in 3’ UTR/ sfRNA1 (**C**). (**D)** Bars represent arithmetic means ± s.e.m. N salivary glands, 43; N saliva, 71; from two mosquito batches. (**E**) Sequences of sfRNA species found in saliva.

Extracellular ncRNAs can be packaged in extracellular vesicles (EVs), including microvesicles and exosomes (Kołat et al., 2019), or in ribonucleoprotein complexes with RNA binding proteins (Maori et al., 2019). Therefore, to determine how sfRNA is secreted and protected in saliva, we subjected infected saliva to different biochemical treatments and tested their effect on sfRNA levels. We used *in vitro* synthesized sfRNA1, which we added to uninfected mosquito saliva, as a control for these experiments. sfRNA1 incubated in uninfected saliva was relatively stable, but as expected this RNA could be digested by adding RNases A/T1 (Fig. 2A). In contrast, the sfRNA1 found in infected saliva was resistant to RNase treatment (Fig. 2B). If the infected saliva was pre-treated with Triton X-100, however, the great majority of the sfRNA in saliva was digested by RNase (Fig. 2B). gRNA in saliva was also resistant to RNase, unless pre-treated with Triton X-100 (Fig. 2C), presumably because it is packaged in virions. Our assays indicated that saliva sfRNA is present in detergent-sensitive structures that protect it from nuclease digestion.

**Figure 2.**
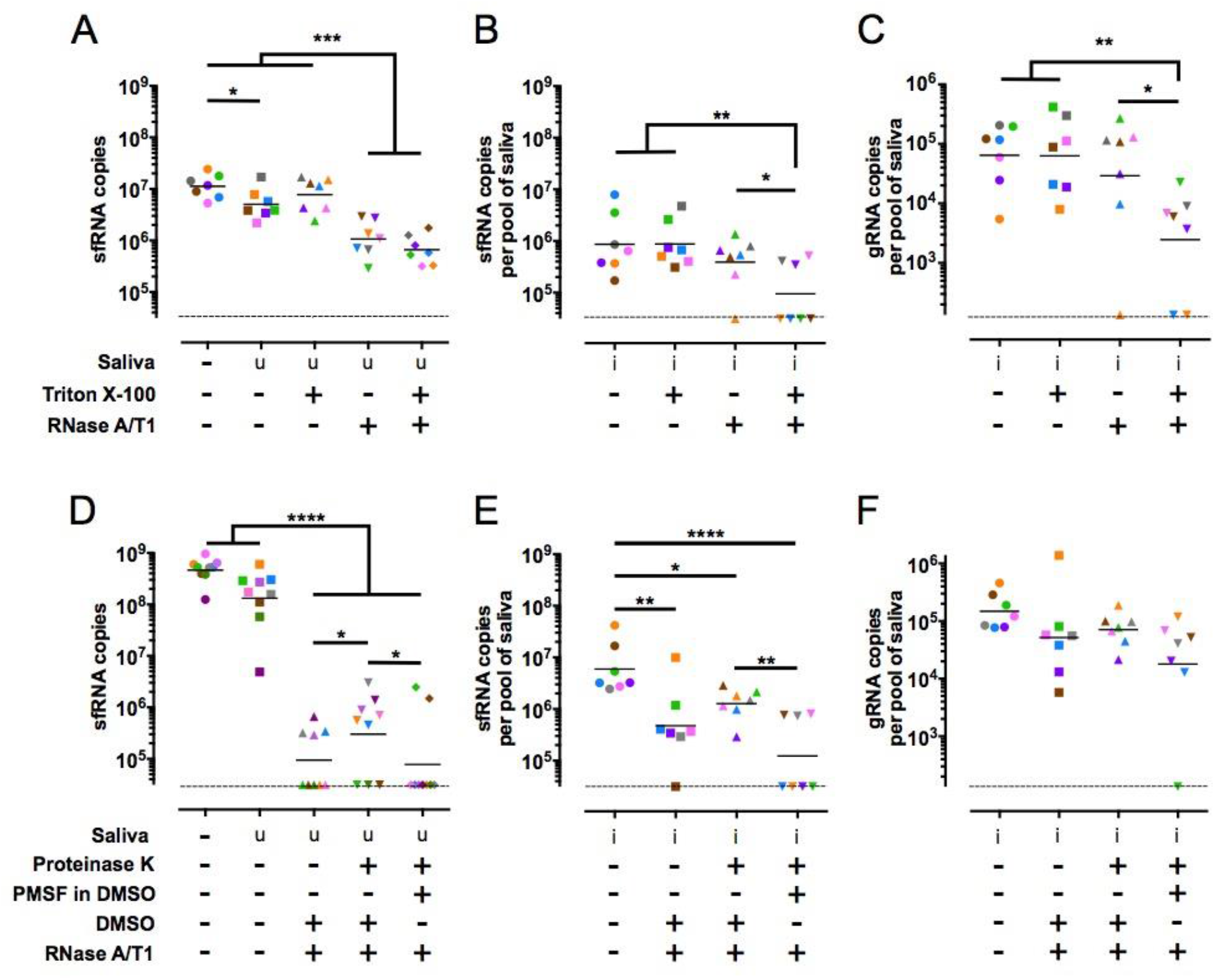
Salivary sfRNA is sensitive to nuclease degradation only after detergent treatment. (**A**-**C**) Impact of Triton X-100 treatment on nuclease resistance for *in vitro* transcribed sfRNA1 in uninfected saliva (**A**) and for sfRNA1 (**B**) and gRNA (**C**) from infected saliva. (**D**-**F**) Impact of DMSO and Proteinase K treatment on nuclease resistance for *in vitro* transcribed sfRNA1 in uninfected saliva (**D**) and for sfRNA1 (**E**) and gRNA (**F**) from infected saliva. Compare the third and fourth group in **E** to note that unless inhibited by PMSF PK would decreased RNase activity. Symbol colors indicate biological repeats. Saliva samples were collected from three mosquito batches. Dashed lines indicate limit of detections which were 133 copies in gRNA and 31,216 copies in 3’ UTR/ sfRNA1. u, uninfected. i, infected. *, p-value < 0.05; **, p-value < 0.01; ***, p-value < 0.001, ****, p-value <0.0001 as determined by Fisher’s LSD test.

To test whether the aforementioned detergent-sensitive structures were ribonucleoprotein complexes, such as those identified in honeybee jelly (Maori et al., 2019), we performed sequential Proteinase K (PK) and RNase A/T1 treatments in saliva samples. To prevent PK from digesting the nucleases, we inhibited the former with PMSF, which was dissolved in DMSO (see Fig. 2 legend). *In vitro*-transcribed sfRNA1 added to uninfected saliva was digested by RNase regardless of PK treatment (Fig. 2D). In infected saliva, we noted partial digestion of sfRNA by RNase in the absence of PK treatment and this could be attributed to DMSO, which was used to control for PMSF (compare Fig. 2B & 2E) and is known to destabilize some membranes (He et al., 2012). It should be noted that gRNA was not sensitive to RNase regardless of treatment (Fig. 2F). This suggests that gRNA, unlike sfRNA, is packaged in a DMSO-resistant compartment, which is expected for DENV virions (Lok et al., 2012). Importantly, sfRNA in infected saliva was digested by RNase to the same extent whether or not samples were pretreated with PK (Fig. 2E). This result suggested that sfRNA in saliva is not protected from nuclease degradation by proteins.

While there is ample evidence that sfRNA has potent anti-immune action (Slonchak and Khromykh, 2018), we tested how sfRNA levels influenced saliva-mediated infections of human cells. To do this we took advantage of our observations that salivary sfRNA levels varied between independent mosquito infections. We collected pools of saliva from mosquitoes infected with three DENV2 strains PR6452, PR1940 and PR9963, and identified those that had similar levels of gRNA copies but substantially different sfRNA levels and therefore had either low or high sfRNA:gRNA ratios (Table 1). In four independent experiments, one with PR6452, two with PR1940 and one with PR9963, we compared infection of human Huh-7 cells with saliva with low or high sfRNA: gRNA ratios (Table 1). For each comparison between low and high sfRNA: gRNA ratio saliva pools we infected Huh-7 cells with the equivalent number of viral genomes in the same total volume of saliva, which is known to impact infectivity (Wichit et al., 2016). To achieve this we diluted infected saliva samples with saliva from uninfected mosquitoes as described in Table 1. As a measure of infection, we quantified foci of viral genome replication detected by immunofluorescence with an anti-dsRNA antibody (Fig. 3). The geometric mean of replication foci per cell was 34.7 when cells were inoculated with PR6452-infected saliva with a lower sfRNA:gRNA ratio vs 64.1 for saliva with a higher ratio (Table 1 and Fig. 3B, C). Very similar results were obtained with PR1940-infected saliva pools. In repeat 1, lower sfRNA:gRNA ratio resulted in a geometric mean of 48.5 foci per cell, whereas higher ratio produced 78.8 foci per well (Table 1 and Fig. 3D, E). In repeat 2, lower vs higher ratio resulted in 49.2 vs 91.6 foci per cell, respectively (Table 1 and Fig. 3 Sup. 1A,B). This effect was not observed with PR9963-infected saliva pools (Table 1 and Fig. 3 Sup. 1C,D), which were carried out with 5-10 fold lower viral titers. These results suggest that higher level of sfRNA in mosquito saliva significantly enhances DENV2 infection in human cells.

**Table 1.** Higher sfRNA:gRNA ratio in mosquito saliva increases DENV2 infection of human cells. The experiment number, identity of the virus strain, concentration of gRNA an sfRNA, and sfRNA: gRNA ratio for pools of salivas collected from DENV2 infected mosquitoes are shown in the first five columns. The total volume of salivary inoculum (volume of infected saliva (inf.) + volume of uninfected saliva (uninf.)) used to infect Huh-7 cells and the gRNA copies added per well of Huh-7 cells are indicated in columns six and seven. The genometric mean, lower and upper 95% CI of dsRNA foci per cell at 2 days post infection are in column eight.

| Exp. | DENV2 strain | gRNA (copies/μl) | sfRNA (copies/μl) | sfRNA:gRNA ratio | Total volume of saliva added per well of Huh-7 cells. (μl) | gRNA inoculum (copies/well) | dsRNA foci per cell, geometric mean (lower and upper 95% CI) |
| --- | --- | --- | --- | --- | --- | --- | --- |
| 1 | PR6452 | $8 \times 10^2$ | $7.2 \times 10^3$ | 9 | 130<br>(130 inf. + 0 uninf.) | $1 \times 10^5$ | 34.7<br>(26.2; 46.0) |
| | PR6452 | $9 \times 10^2$ | $4.2 \times 10^4$ | 46.7 | 130<br>(115 inf. + 15 uninf.) | $1 \times 10^5$ | 64.1<br>(45.8; 89.8) |
| 2 | PR1940 (#1) | $4.2 \times 10^3$ | $2.9 \times 10^3$ | 6.7 | 185<br>(65.5 inf. + 119.5 uninf.) | $2.8 \times 10^5$ | 48.5<br>(34.7; 67.9) |
| | PR1940 (#1) | $1.5 \times 10^3$ | $4.6 \times 10^4$ | 30.2 | 185<br>(185 inf. + 0 uninf.) | $2.8 \times 10^5$ | 78.8<br>(63.2; 98.3) |
| 3 | PR1940 (#2) | $4 \times 10^3$ | $2.2 \times 10^3$ | 5.6 | 100<br>(65 inf. + 35 uninf.) | $2.6 \times 10^5$ | 49.2<br>(23.6; 102.2) |
| | PR1940 (#2) | $2.6 \times 10^3$ | $4.3 \times 10^4$ | 16.6 | 100<br>(100 inf. + 0 uninf.) | $2.6 \times 10^5$ | 91.6<br>(53.23; 157.7) |
| 4 | PR9963 | $3.4 \times 10^2$ | $6.4 \times 10^3$ | 18.8 | 110<br>(83 inf. + 27 uninf.) | $2.8 \times 10^4$ | 51.0<br>(30.1; 86.5) |
| | PR9963 | $2.6 \times 10^2$ | $5.4 \times 10^4$ | 208 | 110<br>(110 inf. + 0 uninf.) | $2.8 \times 10^4$ | 54.7<br>(33.8; 88.6) |

**Figure 3.**
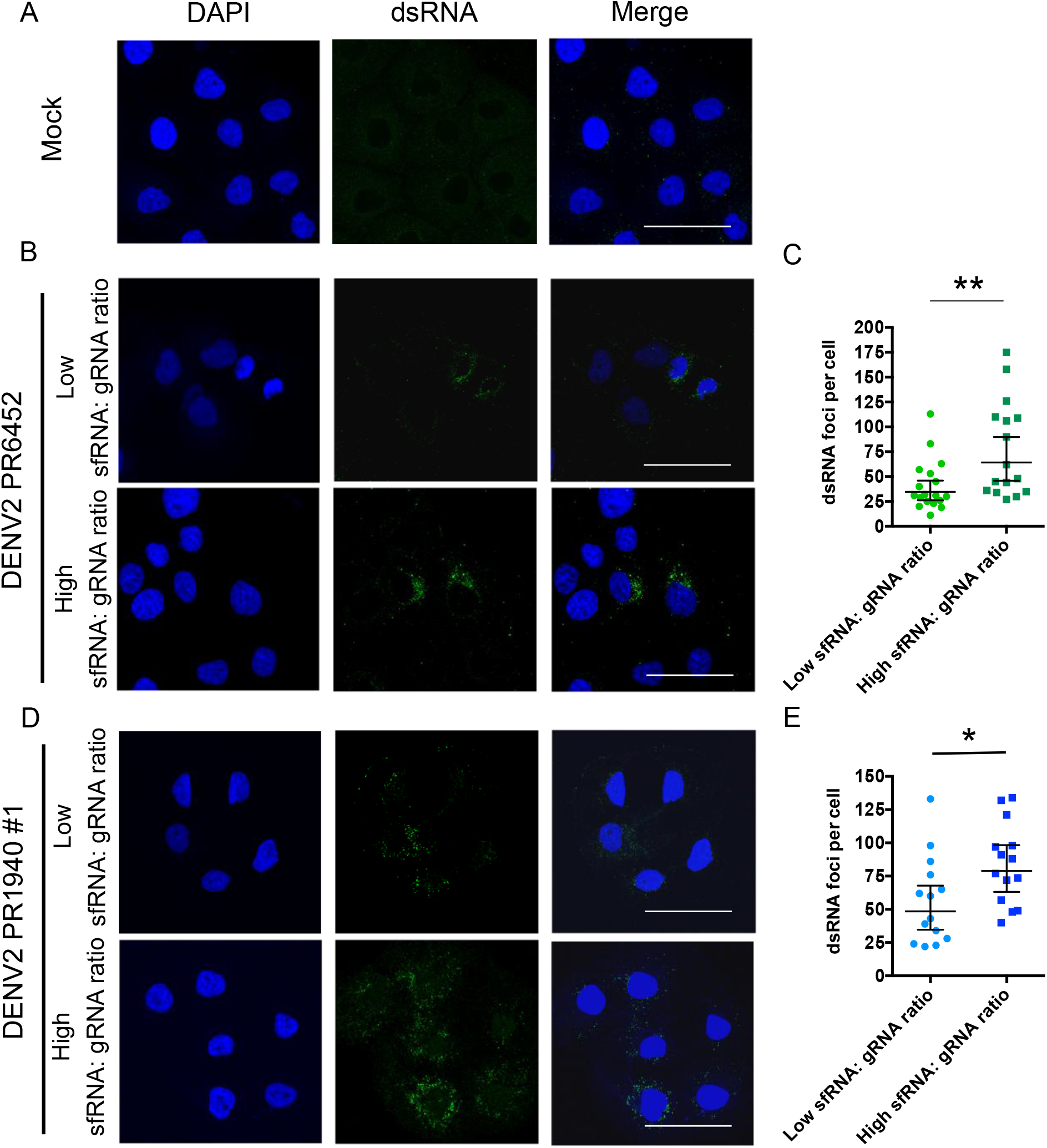
Higher sfRNA:gRNA ratio in mosquito saliva increases DENV2 infection. (**A**) Representative pictures of nucleus (DAPI, blue) and dsRNA (green) from Huh-7 cells supplemented with uninfected saliva. (**B-E**) Representative pictures of nucleus and dsRNA from (B, D) and number of dsRNA foci per (C, E) Huh-7 cell supplemented with saliva infected with DENV2 PR6452 or PR1940 containing either high or low sfRNA:gRNA ratio. (B, D) Scale bar, 50 μm. (C, E) Each point indicates dsRNA foci in one cell. Bars show geometric means ± 95% C.I. *, p-value < 0.05; **, p-value < 0.01 as determined by unpaired t-test. The online version of this article includes the following figure supplement(s) for figure 3: Figure 3, supplement 1. Additional repeat for Huh-7 cells supplemented with PR1940- and PR9963-infected saliva.

Taken together, our results indicate that infection-enhancing sfRNA in infected saliva is located in the lumen of detergent- and DMSO-sensitive vesicles. While it is formally possible that sfRNA is packaged in DENV2 virions in the saliva, as proposed in a recent study (Syenina et al., 2020), we do not favor this conclusion based on the differential DMSO sensitivity observed for gRNA and sfRNA, and our previous data (Bidet et al., 2014). EVs are well known vehicles of coding and noncoding RNAs, including viral RNAs (Nolte et al., 2016). In fact, EVs derived from DENV infected mosquito cells have been isolated, characterized and shown to contain DENV gRNA fragments (Reyes-Ruiz et al., 2019; Vora et al., 2018). Furthermore, EVs derived from arthropod cells in culture have been shown to transmit viral genomes to human keratinocyte-like HaCaT cells (Zhou et al., 2018). Based on these considerations, we propose that sfRNA in saliva is present in EVs, which can deliver their cargo to cells in the same location and at the same time as virus is deposited in the biting site (Fig. 4). sfRNA has been shown to have potent anti-immune function in human cells (Bidet and Garcia-Blanco, 2014; Slonchak and Khromykh, 2018) and here we show that higher levels of sfRNA in saliva correlate with enhanced DENV2 infectivity; therefore, we posit that sfRNA delivered to human skin cells during biting will dampen the local innate immune response and give the virus an advantage that favors mosquito transmission.

**Figure 4.**
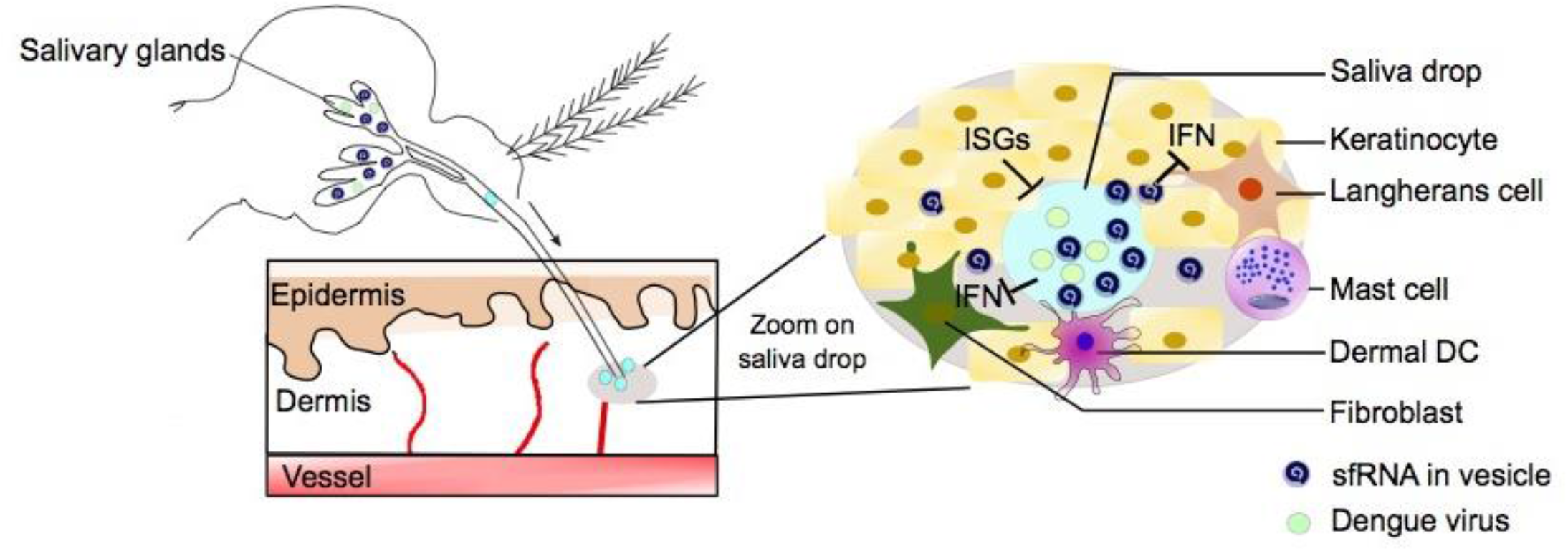
Model: sfRNA in salivary vesicles is transported to cells in the vicinity of first infections at the biting site. sfRNA found in detergent-sensitive vesicles in infectious mosquito saliva is transferred to human skin cells during biting. The anti-immune properties of sfRNA inhibit the antiviral early innate immune response (interferon (IFN) and interferon stimulated genes (ISGs)) to enhance skin cell virus infection and transmission.

The role of EVs in the battle between pathogens and hosts is not unprecedented. The nematode *Heligmosomoides polygyrus* uses EVs to deliver miRNAs and other transactivators to suppress Type 2 innate immunity (Buck et al., 2014). Similarly, hosts can transfer small RNAs to silence virulence genes of pathogens (Cai et al., 2018). In this report, we propose that a viral pathogen harnesses the biology of its invertebrate vector to increase transmission to a human host.

## Materials and methods

### Cell lines, virus, and mosquitoes

*Aedes albopictus* C6/36 (CRL-1660) and baby hamster kidney BHK-21 (CCL-10) cell lines from ATCC were grown in RPMI media (ThermoFisher Scientific), supplemented with 10 % heat-inactivated fetal bovine serum (FBS) (ThermoFisher Scientific) and 1 % Penicillin-Streptomycin (ThermoFisher Scientific) at 28 °C and 37 °C, respectively, in 5 % CO_2_. Huh-7 cell line was maintained in Dulbecco’s modified Eagle medium (DMEM, ThermoFisher Scientific), supplemented with 10 % FBS and 1 % Penicillin-Streptomycin at 37 °C in 5 % CO_2_. Dengue virus 2 (DENV2) New Guinea C (NGC) strain from ATCC (VR-1584) was propagated in C6/36 and titrated using plaque assay in BHK. Low-passage stocks of DENV2 PR1940, PR6452, and PR9963 strains (Manokaran et al., 2015) were gifts from the Dengue Branch of the Centers for Desease Control and Prevention, San Juan, Puerto Rico, and were obtained from Prof. Eng Eong Ooi’s laboratory at Duke-NUS medical School. The *Aedes aegypti* colony was established and reared as detailed previously (Pompon et al., 2017).

### Mosquito oral infection

Three- to five-day-old female mosquitoes were starved for 17 h and offered a blood meal containing 40 % volume of washed erythrocytes from specific-pathogen-free pig’s blood (PWG Genetics), 5 % of 10 mM ATP (ThermoFisher Scientific), 5 % human serum (SigmaAldrich), 50 % of RPMI media and 10^7^ plaque forming unit (pfu) per ml in all experiments. Blood meal titer was validated using plaque assay. Mosquitoes were exposed to the blood meal for 1.5 h and fully engorged females were selected and maintained with water and 10 % sugar solution until analysis at 10 days and 15 days post blood meal in NGC and PR strains, respectively.

### Northern blots

Northern blots were conducted using NorthernMax Kit (Ambion) with modifications to manufacturer’s protocol as described (Bischoff et al., 2004; Filomatori et al., 2017; Pompon et al., 2017). Total RNA from 50 orally-infected mosquitoes was extracted using RNAzol RT (Molecular Research Center) and separated on a denaturing gel with 5 % acrylamide (Bio-Rad) and 8 M Urea (1^st^ Base). Biotinylated single-stranded RNA ladder (Kerafast) was also loaded on the gel. RNA was transferred onto a nylon membrane (Hybond-N; Biodyne B) using Trans-Blot Turbo (Bio-Rad) at 25 V for 30 min. The membrane was UV-crosslinked, pre-hybridized and hybridized with a biotin-16-dUTP (Roche) labeled dsDNA probe including the whole 3’UTR and generated using detailed primers (Bidet et al., 2014). After washes and blocking, the membrane was stained with IRDYE 800cw streptavidin (LI-COR). Pictures were taken with Odyssey CLx imaging system (LI-COR).

### Mosquito saliva collection

Starved mosquitoes were immobilized by cutting wings and legs. Mosquito proboscises were individually inserted into 20 µl tips containing 10 µl of pre-warmed RPMI media with 0.5 mM ATP and 0.01 % fluorescent Rhodamine (Sigma) for 30 min. RPMI mixtures from mosquitoes that ingested Rhodamine, hence salivated, were collected.

### Absolute quantification of gRNA and sfRNA1 copies by real-time RT-qPCR

Mosquito salivary glands were homogenized with 1.0 mm silica beads (BioSpec) using mini-beadbeater-96 (BioSpec) in 350 µl of TRK lysis buffer and saliva-containing mixtures were lysed in 350 µl TRK lysis buffer. Total RNA was extracted with E.Z.N.A. Total RNA kit I (Omega Bio-Tek). gRNA and 3’UTR/ sfRNA1 were quantified in absolute numbers using one-step RT-qPCR (Supplementary Table 1) and standard curves as detailed (Pompon et al., 2017).

To calculate the limit of detection (LoD), gRNA and 3’UTR/ sfRNA1 fragments encompassing the qPCR targets and generated for the standard curves were 10- or 2-fold serially diluted from 1.2 × 10^7^ to 1.9 copies for gRNA and from 6 × 10^8^ to 117 copies for sfRNA1. Four technical repeats per dilution of three independent repeats were quantified using one-step RT-qPCR. Fractions of positive replicates per copies of gRNA or 3’UTR/ sfRNA1 were plotted against a sigmoid curve to identify LoD at 95% confidence (Forootan et al., 2017).

sfRNA1 copy number was calculated by subtracting the number of sfRNA1 and 3’UTR fragments (estimated using the 3’UTR/ sfRNA1 primers) to the number of gRNA fragments (estimated using the envelope primers). For infected samples that contained detectable amount of sfRNA, sfRNA: gRNA ratio was calculated by dividing the number of sfRNA over the number of gRNA. gRNA detection rate corresponded to the number of samples with detectable amount of gRNA over the total number of analyzed samples. sfRNA detection rate corresponded to the number of samples with detectable amount of sfRNA over the number of samples with detectable amount of gRNA.

### sfRNA sequencing

Total RNA from 50 whole mosquitoes or 10 saliva collected after oral infection was extracted using RNAzol RT. sfRNA fragments were circularized using T4 RNA ligase 1 (New England Biolabs) at 4 °C overnight (Filomatori et al., 2017), reverse transcribed using SuperScript III (ThermoFisher Scientific) with primer 5’-GCTGTTTTTTGTTTCGGG-3’ located at nucleotides 10621-10604 of DENV2 NGC genome and amplified using forward primer 5’-AAAATGGAATGGTGCTGTTG-3’ (nucleotides 10690-10709 from NGC genome) and the primer used for reverse transcription (Filomatori et al., 2017). PCR products were gel-separated and the expected-size bands were cloned with Topo TA kit (ThermoFisher Scientific) before sequencing (1^st^ Base).

### RNase resistance assay after detergent or proteinase treatments

Twenty saliva collected from orally-infected or uninfected mosquitoes were pooled and divided into four equal volume. Each subset was used to test one of the four combinations of RNase with either Triton X-100 or proteinase K (PK) treatments. *In vitro* transcribed sfRNA1 generated for standard curve (see above) was added to each uninfected saliva subgroup. Samples were treated with 0.1 % Triton X-100 (Sigma) at room temperature for 30 min before adding 3 µl of RNase A/T1 (ThermoFisher Scientific) at 37 °C for 30 min. Samples were treated with 0.25 µg of PK (ThermoFisher Scientific) at room temperature for 30 min and with 2 mM phenylmethylsulfonyl fluoride (PMSF) (Sigma) at room temperature for 10 min to inhibit PK before adding 3 µl of RNase A/T1 at 37 °C for 30 min. As PMSF was dissolved in DMSO (Sigma), 1% DMSO was added as control when PMSF was not added. RNA was extracted using QIAamp viral RNA mini kit (Qiagen) and sfRNA and gRNA were quantified.

### Infection of human cells by incubation with infected mosquito saliva

3,000 Huh-7 cells were seeded in each well of a 12-well removable chamber slide (ibidi GmbH) and grown in 200 µl of complete media. Pools of saliva from 40 mosquitoes orally-infected with 10^7^ pfu of DENV2 per ml of blood as detailed above were collected in DMEM medium supplemented with 0.5 mM ATP from independent oral infections. Media was removed from Huh-7 cells and cells were incubated with mosquito saliva pools for two hours at 37°C at which point saliva inoculum was removed and replaced with 200 µl of DMEM supplemented with 2% FBS (See Table 1). Two days post inoculation, cells were fixed with 4% paraformaldehyde (PFA) for 20 min, permeabilized with 0.1 % SDS (Merck) in PBS for 10 min, blocked with 0.1 % of BSA in PBS for 40 min, and incubated with 1:500 anti-dsRNA J2 antibody (Sigma-Aldrich) in blocking solution for 1 h at RT. After two times PBS washing, cells were incubated with 1:1000 Alexa Fluor 488-labeled anti-mouse IgG antibody (Invitrogen) in blocking media for 1 h at RT. After washes, slides were mounted with Prolong gold Antifade Mountant containing DAPI (Invitrogen). Pictures were taken with TCS SP8 STED 3X nanoscope (Leica). dsRNA foci were detected and quantified with ImageJ (Schneider et al., 2012) using an in-house macro (Supplementary Table 2) that processed the background at 50 pixels, adjusted the threshold to remove the noise signal, and quantified particle size larger than 0.08 µm^2^.

### Statistical analysis

Differences in sfRNA:gRNA ratio, log-transformed gRNA and sfRNA copies, and log-transformed dsRNA foci were tested with unpaired student’s t-tests. Multiple comparisons with Fisher’s LSD test were used for resistance assays. Prism v5 (GraphPad) was used for analysis.

## Acknowledgments

We thank Drs. Shelton Bradrick, Robert Tesh, Scott Weaver, and Nikos Vasilakis at UTMB, and Drs. Subhash Vasudevan and Eng Eong Ooi at Duke-NUS Medical School for insightful comments. We also thank members of the Garcia-Blanco-Bradrick Laboratory (UTMB) and Garcia-Blanco-Pompon Laboratory (Duke-NUS) for their important suggestions and support. We are also thankful to Drs. Patrick Casey and Wang Lin-Fa, and Ms. Milly Ming Ju Choy, for their support of the Programme insectary and Menchie Manuel for mosquitoes culturing. Finally, we are very grateful to Dr. Lee Ching Ng and her colleagues at National Environment Agency for access to their insectary facility.

## Author contributions

Conceptualization: MAGB and JP.

Formal analysis: SCY, MAGB and JP.

Funding acquisition: RMK, MAGB and JP.

Data acquisition: SCY, WLT, VC, AC

Project administration: MAGB and JP.

Visualization: SCY, MAGB and JP.

Writing – original draft: SCY, WLT, MAGB and JP.

Writing – review and editing: SCY, WLT, VC, AC, RMK, MAGB and JP.

## Data and material availability

All data are available in the main text or the supplementary materials.

## Figures and Tables

**Supplementary Table 1.**
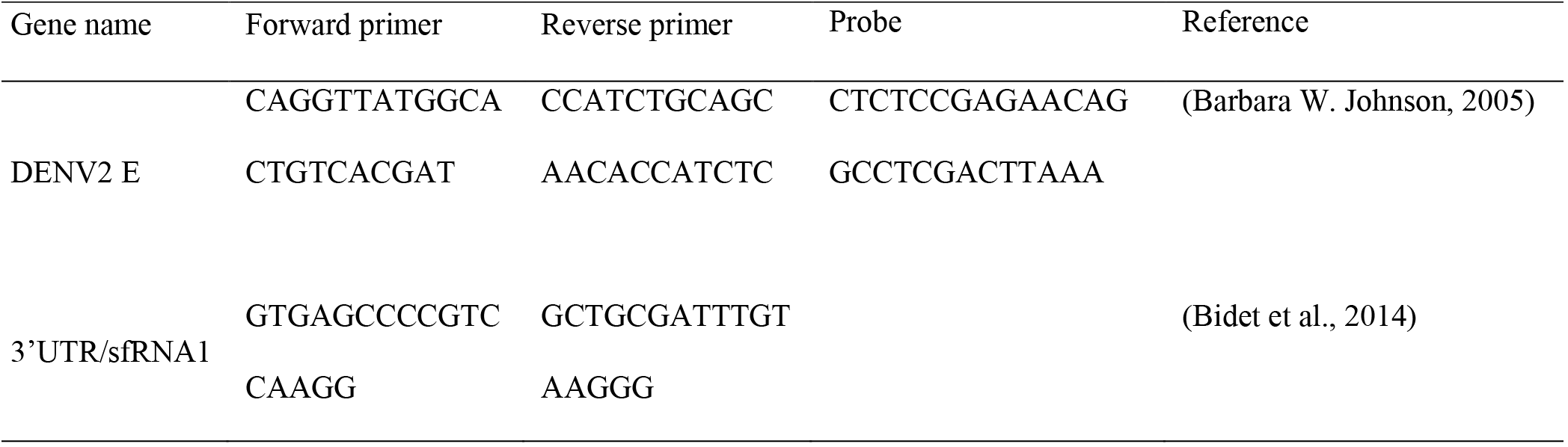
Primer sets used for Real-Time qPCR

| Supplementary Table 1. Primer sets used for Real-Time qPCR |  |  |  |  |
| --- | --- | --- | --- | --- |
| Gene name | Forward primer | Reverse primer | Probe | Reference |
| DENV2 E | CAGGTTATGGCA | CCATCTGCAGC | CTCTCCGAGAACAG | (Barbara W. Johnson, 2005) |
|  | CTGTCACGAT | AACACCATCTC | GCCTCGACTTAAA |  |
| 3'UTR/sfRNA1 | GTGAGCCCCGTC | GCTGCGATTTGT |  | (Bidet et al., 2014) |
|  | CAAGG | AAGGG |  |  |

**Supplementary Table 2.**
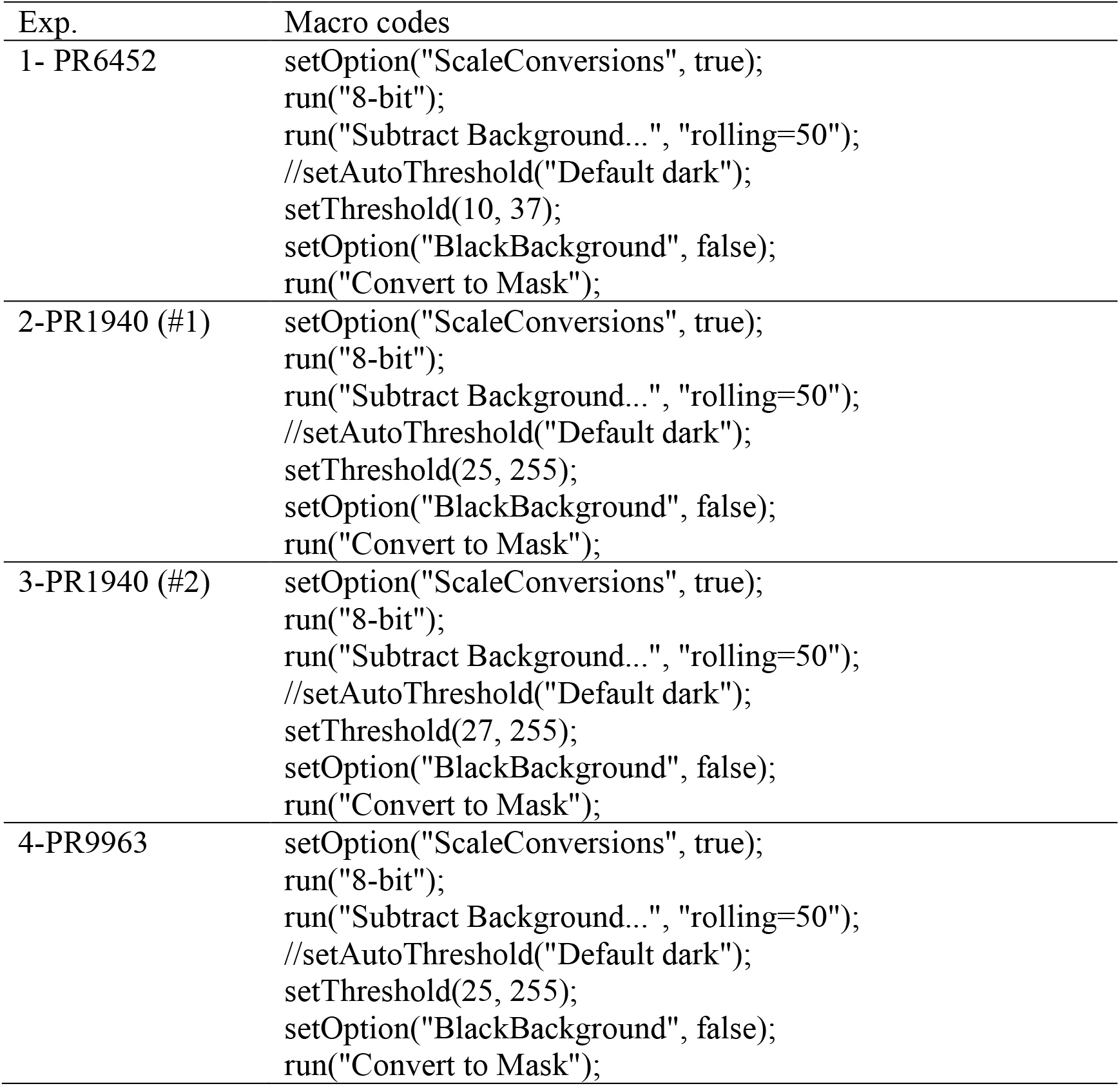
The macro codes for dsRNA foci detection in Image J

| Exp. | Macro codes |
| --- | --- |
| 1- PR6452 | setOption("ScaleConversions", true);<br>run("8-bit");<br>run("Subtract Background...", "rolling=50");<br>//setAutoThreshold("Default dark");<br>setThreshold(10, 37);<br>setOption("BlackBackground", false);<br>run("Convert to Mask"); |
| 2-PR1940 (#1) | setOption("ScaleConversions", true);<br>run("8-bit");<br>run("Subtract Background...", "rolling=50");<br>//setAutoThreshold("Default dark");<br>setThreshold(25, 255);<br>setOption("BlackBackground", false);<br>run("Convert to Mask"); |
| 3-PR1940 (#2) | setOption("ScaleConversions", true);<br>run("8-bit");<br>run("Subtract Background...", "rolling=50");<br>//setAutoThreshold("Default dark");<br>setThreshold(27, 255);<br>setOption("BlackBackground", false);<br>run("Convert to Mask"); |
| 4-PR9963 | setOption("ScaleConversions", true);<br>run("8-bit");<br>run("Subtract Background...", "rolling=50");<br>//setAutoThreshold("Default dark");<br>setThreshold(25, 255);<br>setOption("BlackBackground", false);<br>run("Convert to Mask"); |

**Figure 1, supplement 1.**
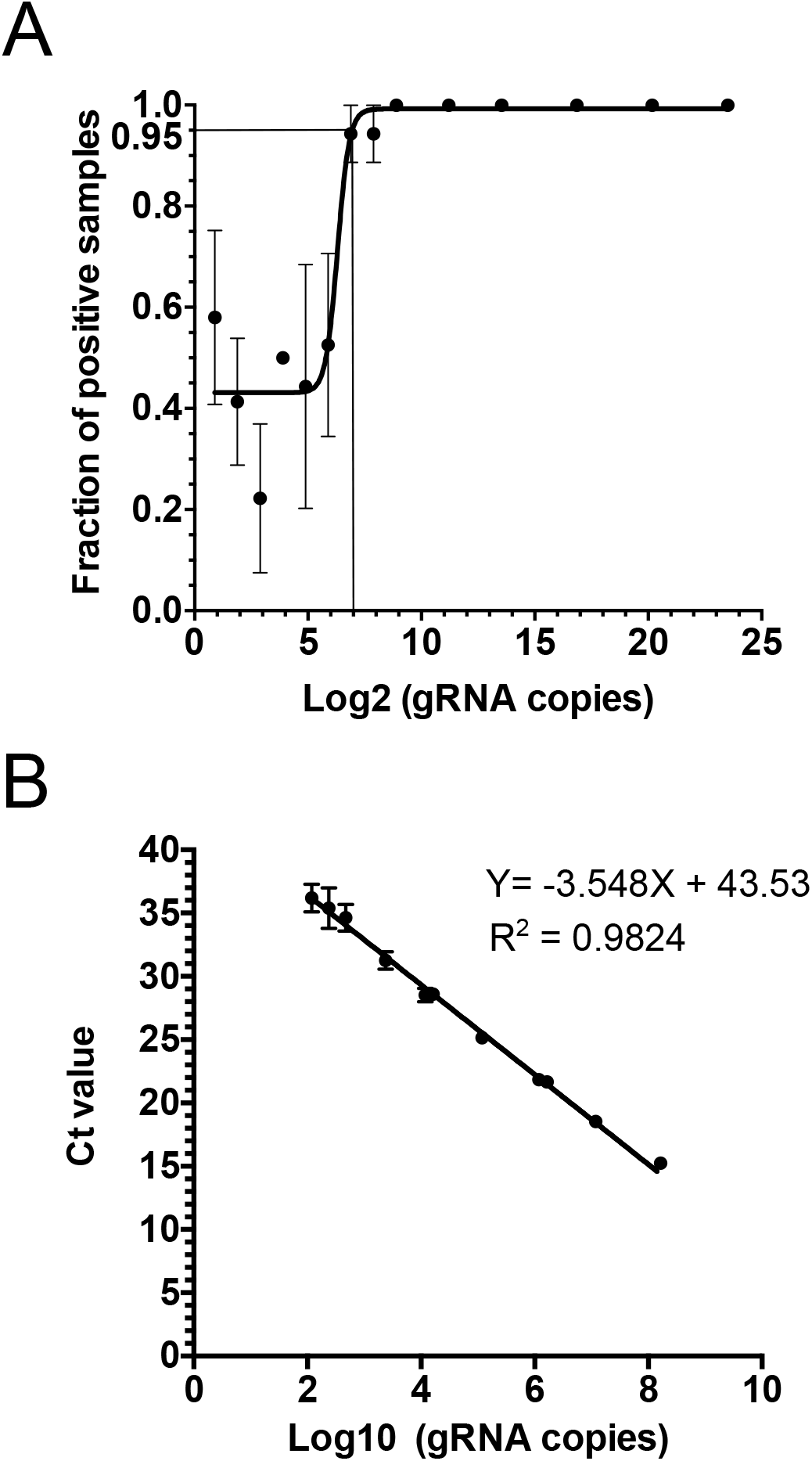
Limit of detection (LoD) and equation for absolute quantification of DENV2 gRNA. *In vitro* transcribed RT-qPCR target was serially diluted from 1.2 × 10^7^ to 1.9 copies before quantification by RT-qPCR. (**A**) Fractions of positive replicates were plotted against gRNA copies. At 95% confidence, LoD of log2 DENV2 gRNA is 7.065, equaling 133 copies. (**B**) Standard curve for absolute quantification of gRNA copies. (**A**-**B**) Points show means ± s.e.m. from 12 technical replicates from 3 independent experiments.

**Figure 1-figure supplement 2.**
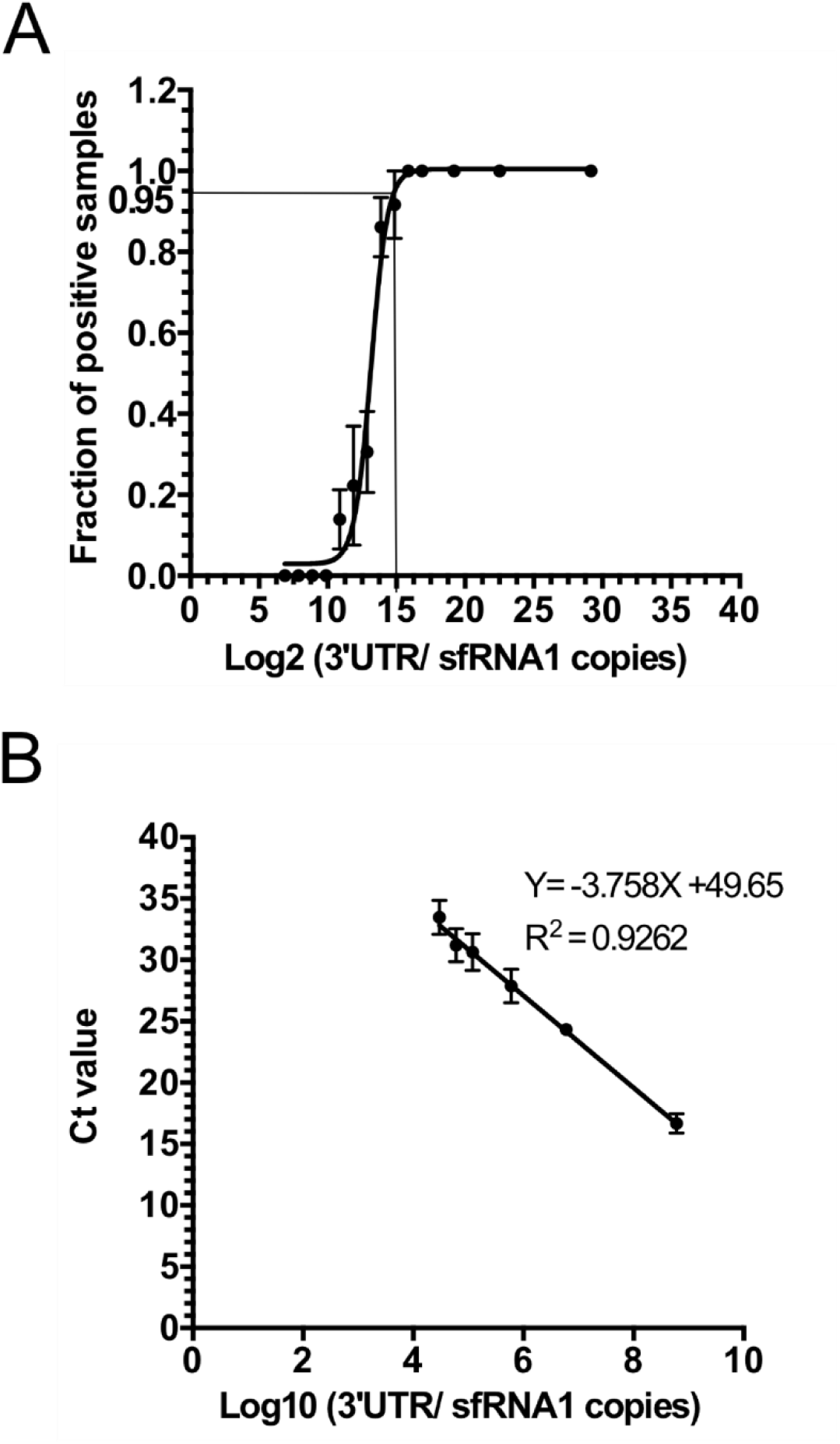
Limit of detection (LoD) and equation for absolute quantification of DENV2 3’ UTR/ sfRNA1. *In vitro* transcribed RT-qPCR target was serially diluted from 6 × 10^8^ to 117 copies before quantification by RT-qPCR. (**A**) Fractions of positive replicates were plotted against sfRNA copies. At 95% confidence, LoD of log2 DENV2 3’UTR/ sfRNA1 is 14.93, equaling 31,216 copies. (**B**) Standard curve for absolute quantification of 3’UTR/ sfRNA1 copies. (**A**-**B)** Points show means ± s.e.m. from 12 technical replicates from 3 independent experiments.

**Figure 1, supplement 3.**
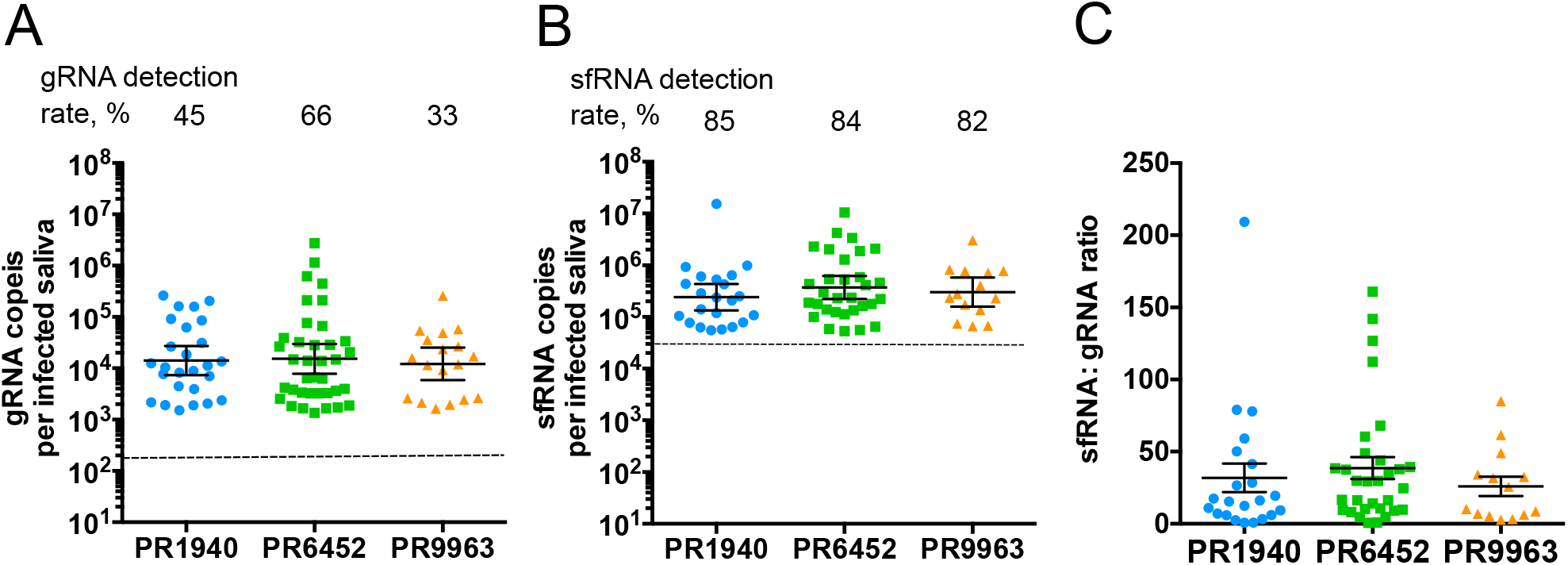
Detection of sfRNA in DENV2 PR strains infected mosquito saliva. (**A**) gRNA; (**B**) sfRNA; and (**C**) sfRNA: gRNA ratio in DENV2 PR1940, PR6452, and PR9963 infected saliva. Points represent one saliva collected from one mosquito. (**A**-**B**) Bars represent geometric means ± 95% C.I. and dashed lines indicate limit of detections which were 133 copies in gRNA (**A**) and 31,216 copies in 3’ UTR/ sfRNA1 (**B**). (**C)** Bars represent arithmetic means ± s.e.m. N PR1940 saliva, 58; N PR6452 saliva, 56; N PR9963 saliva, 52 from two mosquito batches.

**Figure 1-supplement 4.**
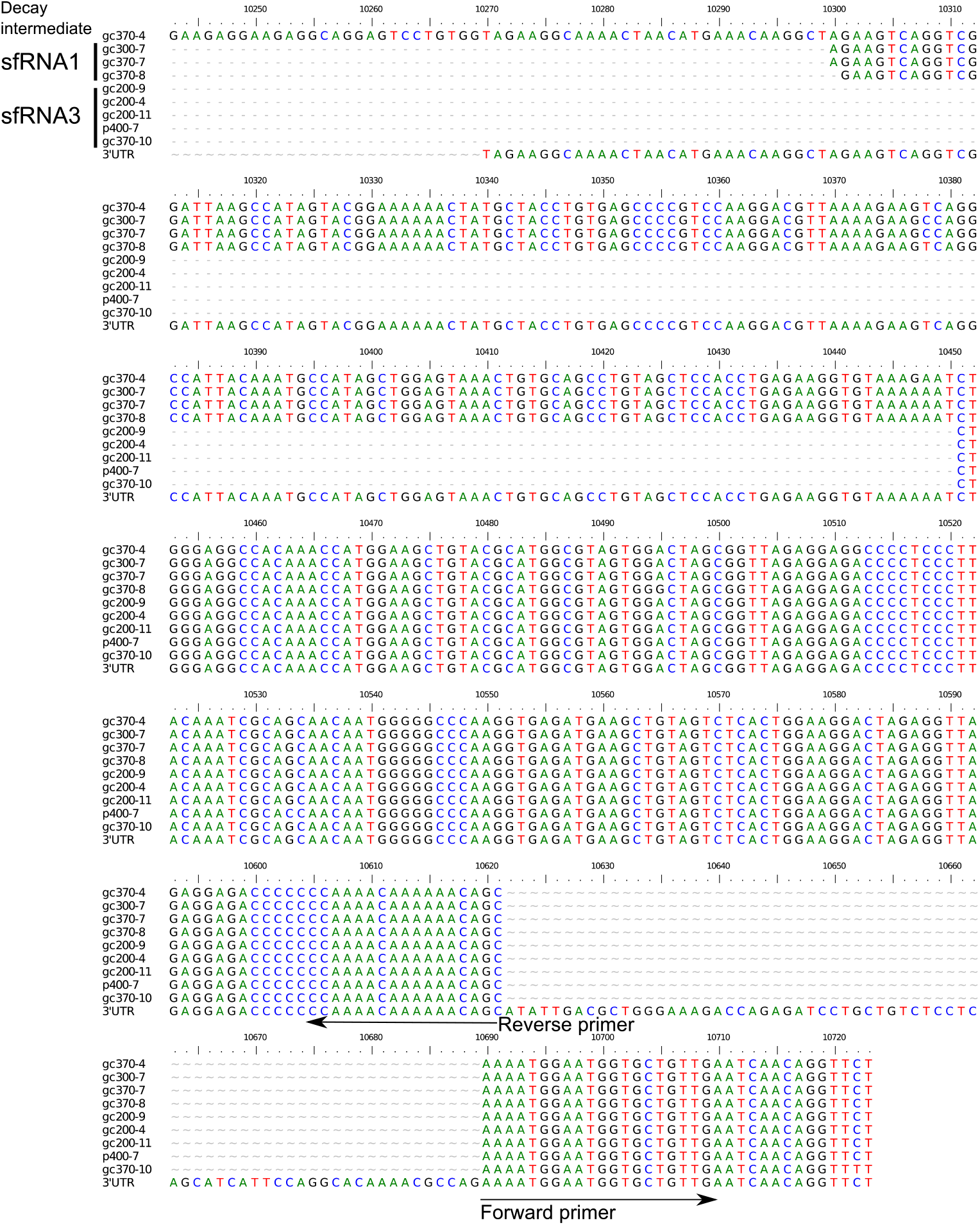
sfRNA sequences from whole mosquitoes. Whole mosquitoes were collected at 10 days post oral infection with DENV2 NGC strain. sfRNA sequences were obtained by RNA circularization, directed expansion of circularized templates, cloning of expected-size amplicons and sequencing. Out of 27 clones sequenced for each sfRNA1 to sfRNA4 expected-sizes, we found sequences corresponding to sfRNA1 and sfRNA3, and one corresponding to a decay intermediate (gc370-4).

**Figure 1-figure supplement 5.**
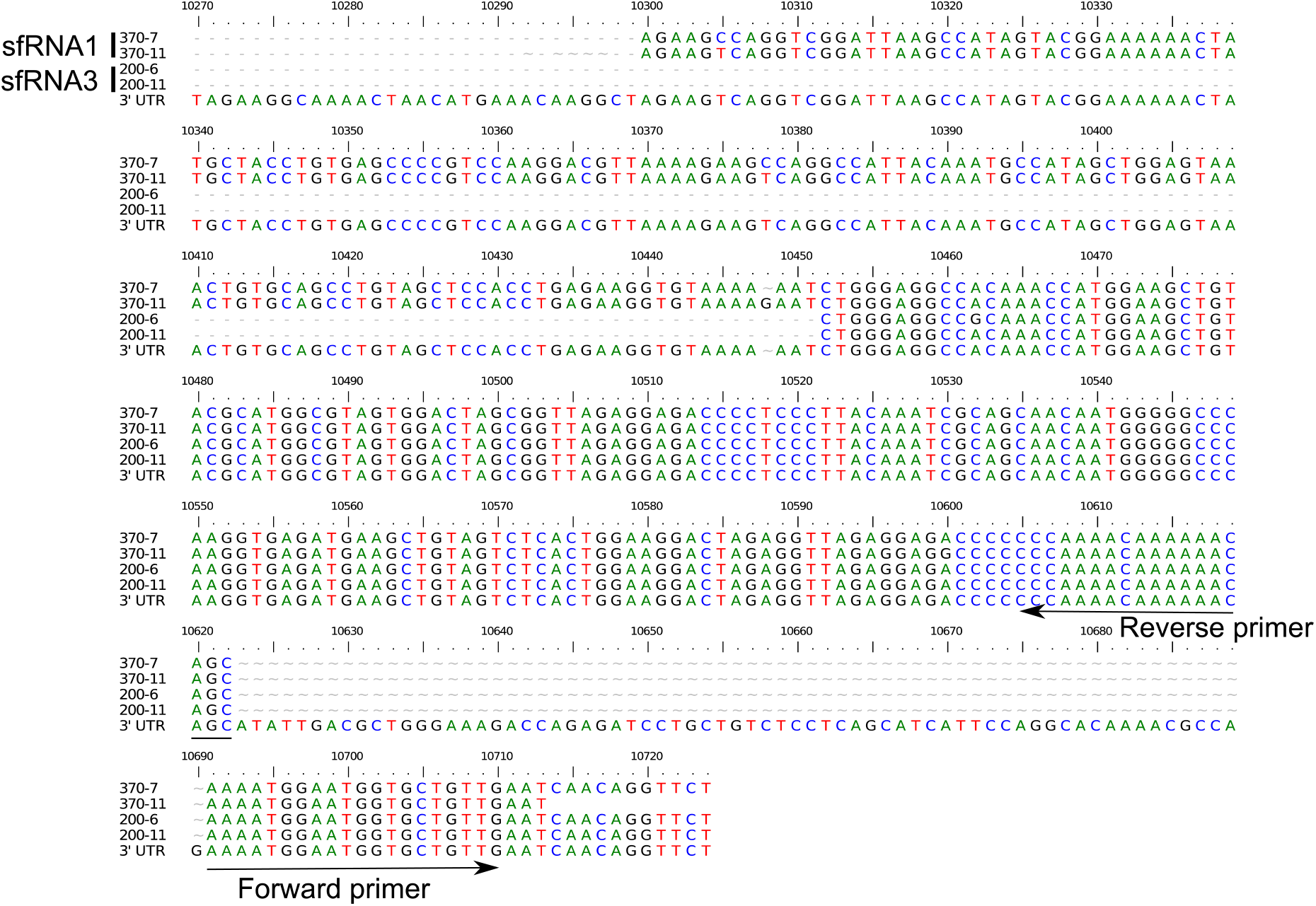
sfRNA sequences from saliva. Saliva samples were collected at 10 days post oral infection with DENV2 NGC strain. sfRNA sequences were obtained by RNA circularization, directed expansion of circularized templates, cloning of expected-size amplicons and sequencing. Out of 26 clones sequenced for each sfRNA1 to sfRNA4 expected-sizes, we found four sequences corresponding to sfRNA1 and sfRNA3.

**Figure 3, supplement 1.**
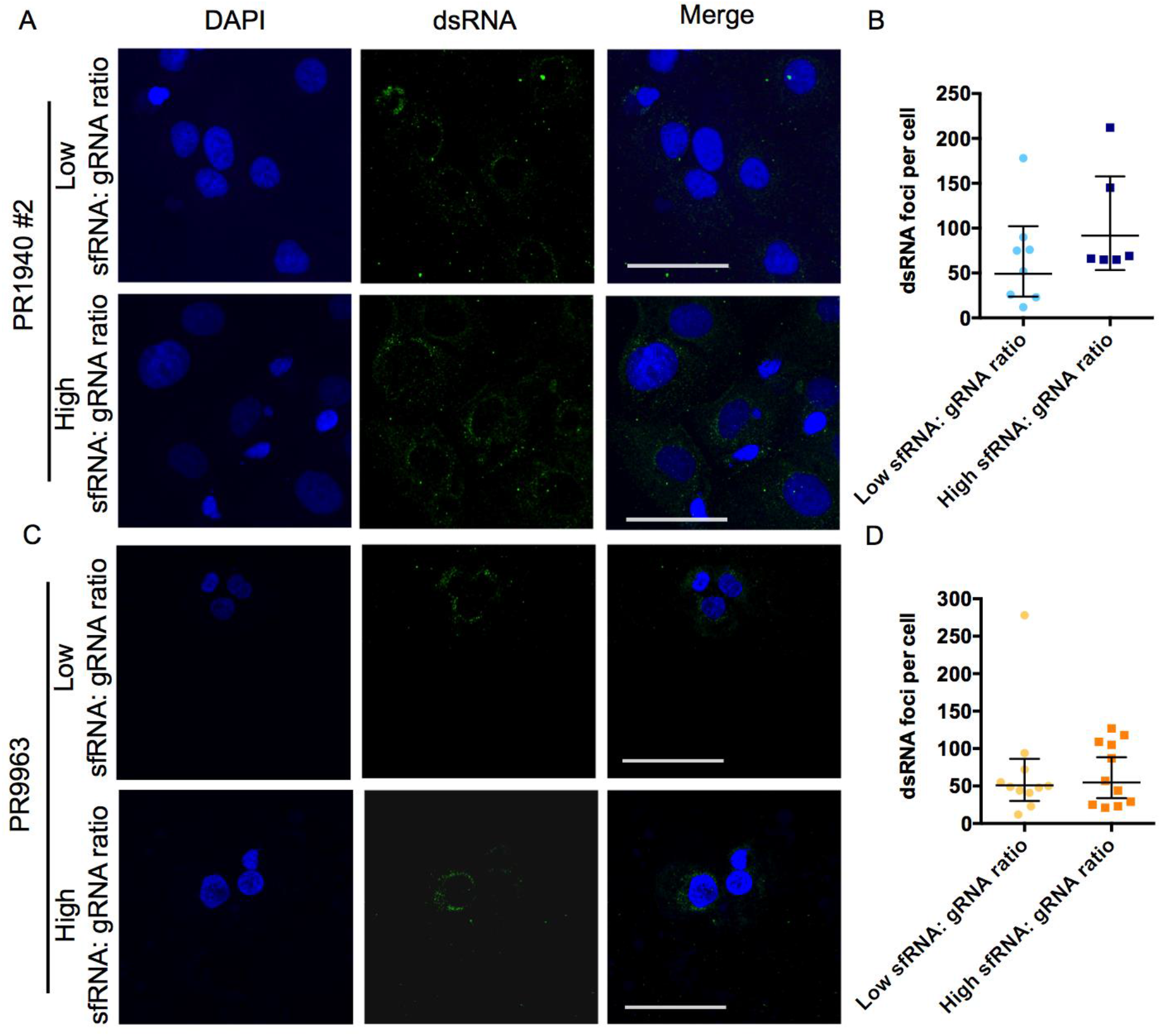
Additional repeat for Huh-7 cells supplemented with PR1940- and PR9963-infected saliva. (**A-D**) Representative pictures of nucleus (DAPI, blue) and dsRNA (green) from (A, C) and number of dsRNA foci per (B, D) Huh-7 cell supplemented with saliva infected with DENV2 PR1940 or PR9963 containing either high or low sfRNA: gRNA ratio. (A, C) Scale bar, 50 μm. (B, D) Each point indicates dsRNA foci in one cell. Bars show geometric means ± 95% C.I..

